# Distinct communities of Bacteria and unicellular eukaryotes in the different water masses of Cretan Passage water column (Eastern Mediterranean Sea)

**DOI:** 10.1101/2024.04.11.588839

**Authors:** Georgia Charalampous, Konstantinos A. Kormas, Eleftheria Antoniou, Nicolas Kalogerakis, Evangelia Gontikaki

## Abstract

Understanding the diversity and dynamics of marine microbiota holds significant importance due to their role in maintaining vital ecosystem functions and services including climate regulation and bioremediation. Here, we studied the diversity and associations between Bacteria and unicellular eukaryotes in the different water masses of the Cretan Passage water column in the Eastern Mediterranean Sea (EMS). Samples were collected from two stations in the Hellenic Exclusive Economic Zone (EEZ) at various depths down to 1000 m during two sampling expeditions in August 2019 and February 2020. Through high-throughput 16S and 18S rRNA gene analysis, we unveiled vertical variations in both bacterial and unicellular eukaryotes diversity respectively. Additionally, interspecies co-occurrence patterns were evaluated between the top and bottom water masses. Our results revealed species fluctuations indicative of seasonality in the surface water mass while the deepest water layers were enriched in heterotrophic taxa and grazers related to organic matter degradation and nutrient cycling. Finally, we found a higher number of microbial associations in surface waters indicating abundant ecological niches compared to the deepest layer, possibly related to the lack of bottom-up resources in the oligotrophic deep ocean.

## Introduction

Deterministic procedures of bottom-up and top-down selection affect the diversity and composition of marine microbial communities and drive the relationships among microorganisms structuring the microbial food web [1, 2]. Despite their importance, the ocean’s microbial biosphere still remains vastly unexplored. However, the advancement of culture-independent methods and the growing DNA and RNA sequence databases, contributed significantly to our knowledge regarding the ecological roles and trophic connections among photo-autotrophic, phago-heterotrophic and mixotrophic marine microbes [3, 4].

The Eastern Mediterranean Sea (EMS), a sub-basin of the semi-enclosed Mediterranean Sea, expands eastwards from the Straits of Sicily and is of great importance both ecologically and geographically. This aquatic system presents complex oceanic processes and is one of the most oligotrophic, saltier and warmer basins of the world [5, 6]. The EMS thermohaline, anti-estuarine circulation leads to stratification into three distinct water masses. Here, the surface Atlantic Water (AW) presents variable salinity (< 38.9 psu) due to evaporation and mixing with the local Levantine Surface Waters (LSW) accumulating in depths down to 150 m. Reaching the eastern-most part of the EMS, the AW/LSW is transformed into the Levantine Intermediate Water (LIW), a more saline (>38.9 psu) and denser water mass, expanding between 150-400 meters in depth and moving in the opposite direction. Finally, the deepest Eastern Mediterranean Deep Water (EMDW) of lower salinity (< 38.9), occupies depths below 400m [5, 7].

One of the most important traits of the EMS is the high deep water temperature (13-15 °C), lower than the overlying water masses but significantly warmer than any other marine system at respective depths, providing a favourable condition for microbial growth and activity [8]. Even though it is a land-locked basin with increasing anthropogenic activity, the EMS waters have been characterised as ultraoligotrophic. Nutrient levels increase with depth having a high nitrate to phosphate ratio (N:P=28:1), exceeding by far the Redfieldian 16:1, making the system P-starved [9]. Furthermore, primary productivity is one of the lowest observed in oceanic systems globally, three times lower than the western Mediterranean and other similar oligotrophic ecosystems (10-143 g C m^-2^ y^-1^) [10].

Under these unique conditions, microbial life thrives, occupying different ecological niches. This low-nutrient and low-primary production environment provides an ecological advantage for mixotrophy, triggering the feeding on microbial prey to sustain carbon and inorganic nutrient budgets and to ensure the functioning of the biological pump and biomass transfer to higher trophic levels [11]. Transmediterranean offshore epipelagic cruises have provided significant insights in terms of longitudinal distribution and analysis for plankton, nanoflagellates and ciliates [12–14]. In the EMS, most studies have been performed in the eastern part of the Levantine basin, recording microbial variations from a temporal, vertical and spatial point of view [15–20]. Whereas studies in the western region of the Levantine Sea, have predominantly used approaches such as flow-cytometry and ^3^H-leucine incorporation methods to determine microbial growth and activity with only a few employing molecular biology tool for the analysis of community composition [21–24]. Nonetheless, microbiome exploration in marine areas off-Crete has been limited [25].

In this study, we explored the microbial community composition across depths, down to 1000 m, in the Cretan Passage water column using the 16S and 18S rRNA gene diversity. We hypothesised that associations between bacteria and unicellular eukaryotes are different between the surface (AW/LSW) and deep (EMDW) water samples and we tested this hypothesis through co-occurrence networks. To our knowledge, no study so far has combined DNA metabarcoding for both bacteria and unicellular eukaryotes to investigate community composition and networking between microorganisms in this marine habitat [15, 17, 23].

## Methods

### Seawater sample collection

Seawater samples were collected in marine areas off South Crete (Cretan Passage) onboard HCMR’s R/V Aegaeo during two oceanographic cruises. Station A (Koufonisi) was sampled in August 2019 and station B (Gavdos) in February 2020 (Fig. 1). Water samples were collected from the surface to 1000 m depth with a SBE911 plus CTD unit, equipped with a 12-bottle SBE32 carousel water sampler. The water density at AW/LSW ranged from approximately 26 to 29 kg m^-3^ for Koufonisi and ∼28.8 kg m^-3^ for Gavdos station respectively while at EMDW the density ranged between 29.1-29.2 kg m^-3^ for both stations (Fig. S1). Temperature, salinity and dissolved oxygen profiles were also built indicating a winter vertical mixing down to 300 m in Gavdos (Fig. S2). Two litres of seawater from each depth were immediately filtered through a 0.2 µM PES membrane filter and were stored frozen in cryotubes for subsequent DNA extraction.

**Fig. 1.**
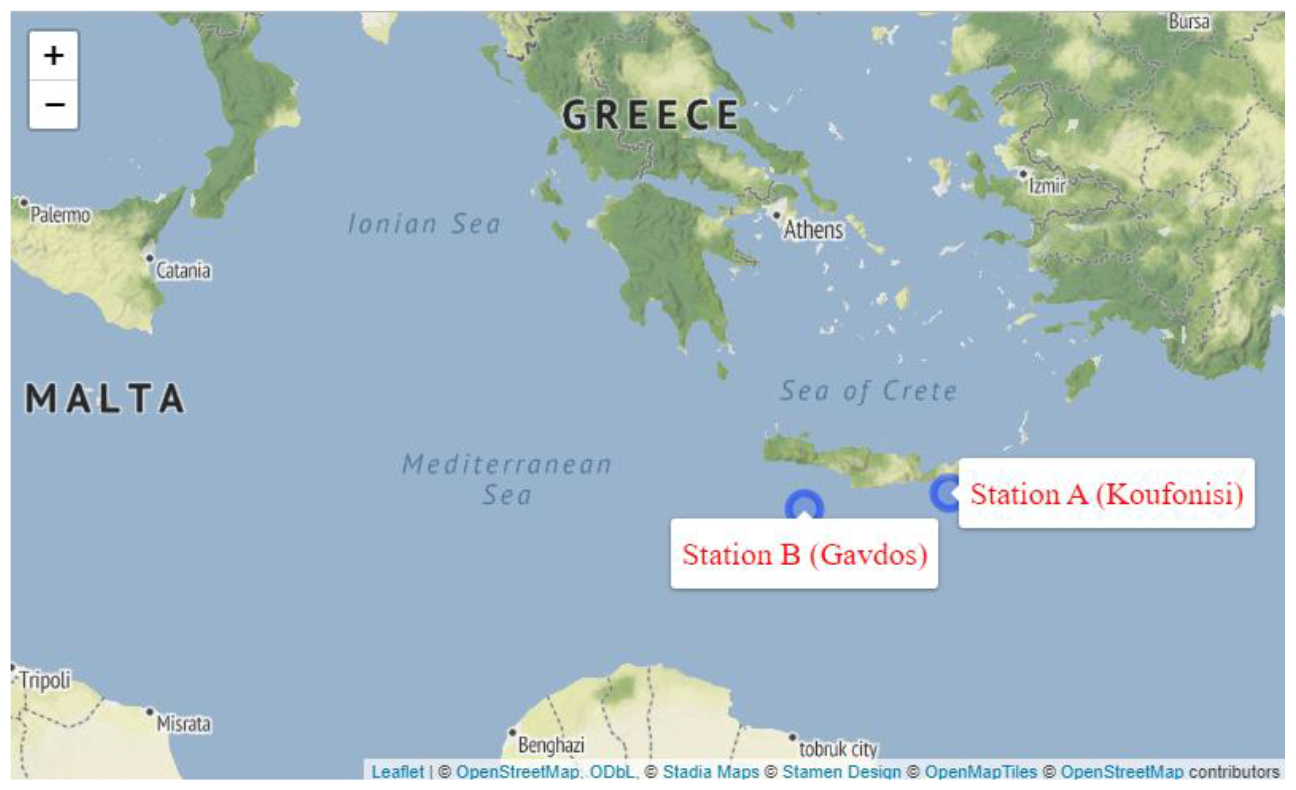
Map pinpointing the two sampling stations

### DNA extraction and sequencing

The method followed for DNA extraction has been previously described in Charalampous et al [26]. In summary, membrane filters were vortexed in CTAB extraction buffer, followed by three freeze-thaw cycles. Subsequently, lysozyme and Proteinase K were added, and the mixture was incubated at 37 °C for 30 min. After treatment with SDS, centrifugation, and extraction with phenol:chloroform:isoamylalcohol, DNA was left to precipitate overnight in isopropanol. The DNA pellet was washed with ethanol, dried, and resuspended in TE buffer. DNA concentration was quantified using Qubit (Invitrogen, Thermo Fisher Scientific, USA) and samples were subjected to high-throughput sequencing. Bacterial and unicellular eukaryotic samples (Table 1) were analysed using the universal primers U341F-U806R for the amplification of the 16S rRNA gene and E572F-E1009R primers for the amplification of the 18S rRNA gene respectively [27, 28]. Sequencing was performed on the Illumina Miseq platform by Biosearch Technologies (LGC Genomics GmbH, Berlin, Germany).

**Table 1.**
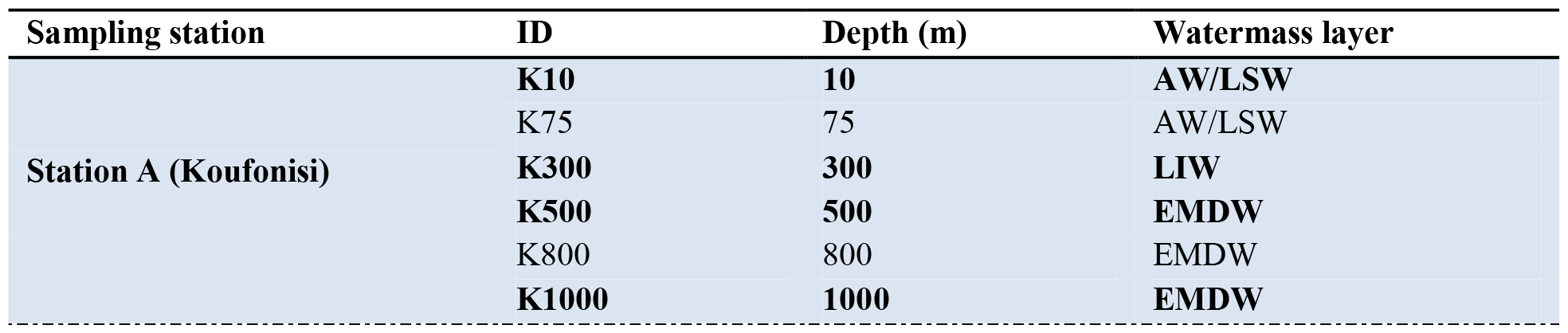
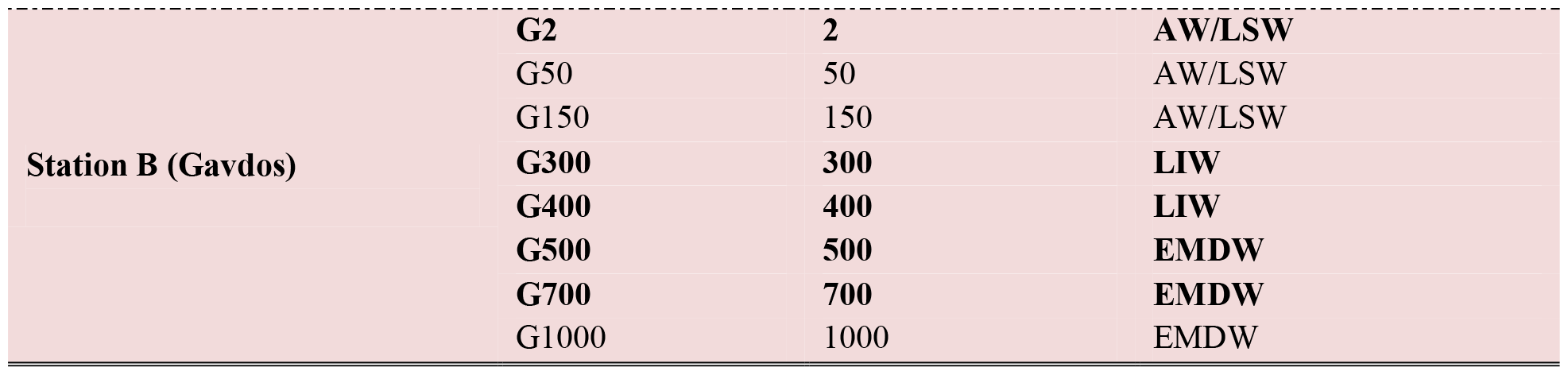
Summary of location, depth and water mass of seawater samples retrieved during the oceanographic expeditions. 16S rRNA sequencing was performed on all samples. In bold are the samples subjected to 18S rRNA sequencing.

| Sampling station | ID | Depth (m) | Watermass layer |
| --- | --- | --- | --- |
| Station A (Koufonisi) | **K10** | **10** | **AW/LSW** |
|  | K75 | 75 | AW/LSW |
|  | **K300** | **300** | **LIW** |
|  | **K500** | **500** | **EMDW** |
|  | K800 | 800 | EMDW |
|  | **K1000** | **1000** | **EMDW** |
| **Station B (Gavdos)** | **G2** | **2** | **AW/LSW** |
|  | G50 | 50 | AW/LSW |
|  | G150 | 150 | AW/LSW |
|  | **G300** | **300** | **LIW** |
|  | **G400** | **400** | **LIW** |
|  | **G500** | **500** | **EMDW** |
|  | **G700** | **700** | **EMDW** |
|  | G1000 | 1000 | EMDW |

### Bioinformatics analysis

DNA sequencing data were analysed using the DADA2 package in Rstudio (version 2022.12.0) [29] in the R programming environment (version 4.2.2) [30]. Bacterial and eukaryotic datasets were created from primer-clipped reads, filtered, and trimmed for consistent length based on quality profiling. Amplicon sequence variants (ASVs) underwent dereplication, denoising, and merging, followed by chimera screening and taxonomy assignment using SILVA SSU (version 138) and PR2 SSU (version 4.14.0) databases for 16S and 18S rRNA sequences, respectively [31, 32]. Two phyloseq objects were constructed for downstream analysis for each microbial domain. Uncharacterised phyla, Chloroplasts, and Archaea were removed from the bacterial dataset, while uncharacterised supergroup taxa, Metazoa, and Streptophyta species were discarded from the eukaryotic dataset. In the end of all downstreaming processes, the bacterial dataset obtained from 14 samples, consisted of 179,556 reads with an average of 12,868 reads per sample (min: 4234, max: 34227) and a total of 2113 ASVs which were distributed among 30 phyla. Whereas, the eukaryotic dataset retained from 9 samples, consisted of 255,508 reads with an average of 28,390 reads per sample (min: 17,305, max: 39,662) and a total of 1678 ASVs were organised into 25 divisions.

### Statistical analysis

Alpha-diversity analysis was performed on ASV abundances prior to normalisation. ANOVA followed by Tukey’s HSD test were used to find statistically significant differences in alpha diversity between water masses and sampling stations (seasonality). Beta-diversity was analysed using PCoA analysis on Bray-Curtis dissimilarity distances after cumulative sum scaling (CSS) normalisation. PERMANOVA (999 permutations) was used to test significant differences using the *adonis2* function of the *vegan* package. ANOVA was also applied to test the significance in relative abundance of bacterial taxa between the surface (AW/LSW) and deep (EMDW) water layers. All figures were generated using the *ggplot* package in R.

### Co-occcurrence network analysis

Co-occurrence network analysis was performed separately for samples collected from surface (AW/LSW) and deep (EMDW) water layers. For each water mass, ASV abundance tables were filtered to keep taxa which were present more than 5 times in at least 50% of the samples for both bacterial and unicellular eukaryotes. Following filtering, the two ASV tables were merged for the correlation process in R. The AW/LSW table consisted of 51 ASVs (31 unicellular eukaryotes, 20 bacteria) and the EMDW consisting of 69 ASVs (57 unicellular eukaryotes, 12 bacteria). Spearman’s correlation was calculated separately for each water mass table using the *Hmisc* package. Networks were constructed in Cytoscape (version 3.9.1) and included only the statistically significant correlations (p<0.05) with R>0.6 or R<-0.6.

## Results

### Beta-diversity

Principal coordinate analysis (PCoA) on Bray-Curtis dissimilarity distances was performed for both bacterial and unicellular eukaryotic datasets indicating that samples were primarily separated according to the water mass from which they were retrieved from (Fig. 2, Fig. S3). AW/LSW samples clustered together while the deepest EMDW samples formed a second group. The microbial communities from the intermediate layer (LIW) clustered together with those from the upper-EMDW depths (G500, K500) besides LIW-G300 sample which grouped along with the AW/LSW samples (Fig. 2, Fig. S3). PERMANOVA analysis indicated statistically significant differences between the microbial communities with water mass (*p*_bacterial_ = 0.001, *p*_eukaryotic_ = 0.017) but not with sampling location (season) (*p*_*bacterial*_ = 0.102, *p*_eukaryotic_ = 0.367). Pairwise PERMANOVA analysis indicated significantly different bacterial communities between the top (AW/LSW) and the bottom (EMDW) water masses (*p*_AW/LSW-EMDW_ = 0.006). Differences between AW/LSW-LIW and LIW-EMDW were less pronounced, yet statistically significant (*p*_AW/LSW-LIW_ = 0.045, *p* _LIW-EMDW_ = 0.033).

**Fig. 2.**
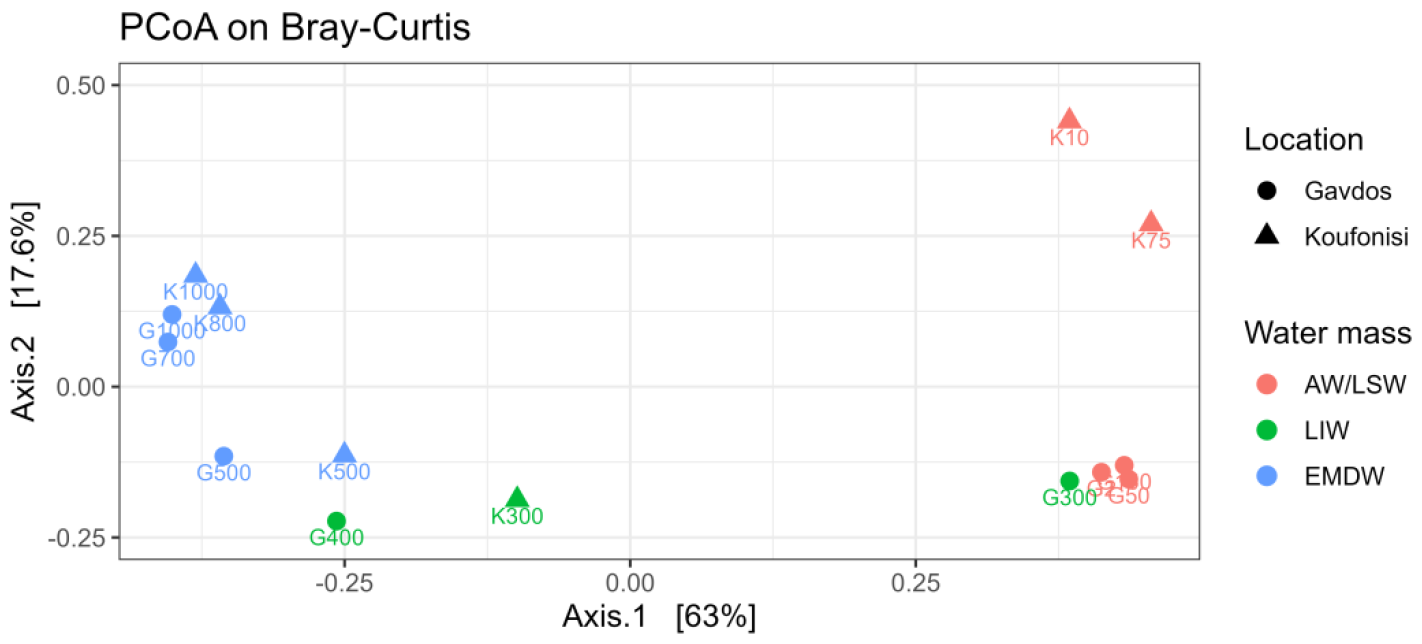
Principal coordinate analysis of bacterial samples using the Bray-Curtis dissimilarity distances

### Alpha-diversity

Bacterial and unicellular eukaryotic diversity were assessed using Shannon, Simpson, and Phylogenetic Diversity (PD) indices. Bacterial analysis revealed lower diversity in the AW/LSW layer compared to LIW and EMDW, with the latter having slightly higher diversity (Fig. 3a, Fig. S4). ANOVA, applied to each metric, revealed only Shannon and PD metrics having a significant difference with water mass (*p*< 0.05). Tukey HSD post-hoc test indicated that AW/LSW differed significantly from LIW and EMDW (Table S1). Moreover, sampling location (season) had no effect on the overall bacterial diversity in contrast to unicellular eukaryotic communities where ANOVA identified significant differences in Shannon and Simpson metrics (Fig. 3b-d, Table S2). Further analysis of eukaryotes showed that alpha diversity was higher in samples from Gavdos (winter) across all three water layers, with the intermediate water mass (LIW) exhibiting the greatest difference compared to Koufonisi. (Fig. S5). Finally, significant eukaryotic diversity between water masses was observed only in Simpson index (Table S2).

**Fig. 3.**
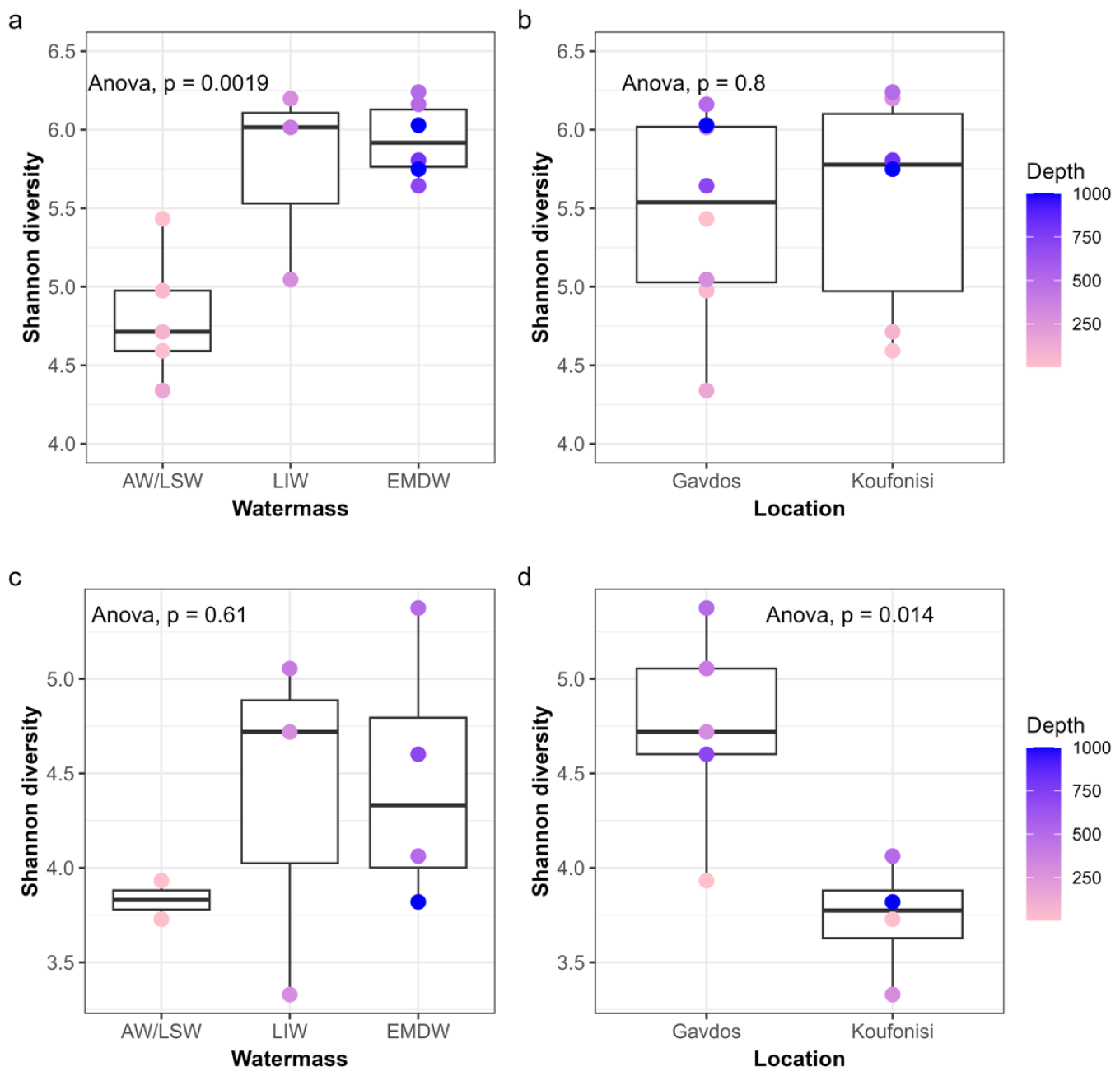
Boxplot of Shannon diversity for bacterial (a-b) and unicellular eukaryotic (c-d) communities in each water mass and sampling location

### Bacterial community structure

Relative abundance analysis revealed water mass-specific bacterial taxa. ANOVA identified 14 phyla with significant differences in abundance between the upper (AW/LSW) and the deepest (EMDW) water layers. Among them were Cyanobacteria (*p*=0.009) and Bacteroidota (*p*=0.012) which were significantly enriched in AW/LSW (Fig. 4). *Synechococcus* dominated Cyanobacteria in AW/LSW, especially in Gavdos (winter), while Bacteroidota were favoured in Koufonisi (summer) comprising 9% of surface ASVs, nearly 20 times higher than EMDW. *Flavobacteriales* were the dominant Bacteroidota in all samples however, in Koufonisi (summer) *Rhodothermales, Balneolales* and *Sphingobacteriales* were also present. Nitrospinota were favoured in intermediate depths (LIW) while Chloroflexi (*SAR202 clade*) were enriched in the deepest water layers (LIW, EMDW), regardless of locality (seasonality) having their highest relative abundance in LIW-G400. Deltaproteobacteria (*SAR324 clade*), Marinimicrobia (*SAR406 clade*), Actinobacteriota (*Microtrichales*) and Nitrospirota were also favoured in bottom water masses and their relative abundance increased with depth (Fig. 4, Fig. S6). For example, Marinimicrobia at 1000m depth comprised approximately the 11% of the bacterial community, twice the average levels present in intermediate depths. Notably, known hydrocarbon degraders (*Alcanivorax, Alteromonas, Pseudoalteromonas, Halomonas, Idiomarina*) were present throughout the water column in Koufonisi (summer) and in deeper waters off Gavdos (winter) station (Fig. S7). Overall, rare taxa (<1% abundance, “Other”) increased with depth in the water column (Fig. 4).

**Fig. 4.**
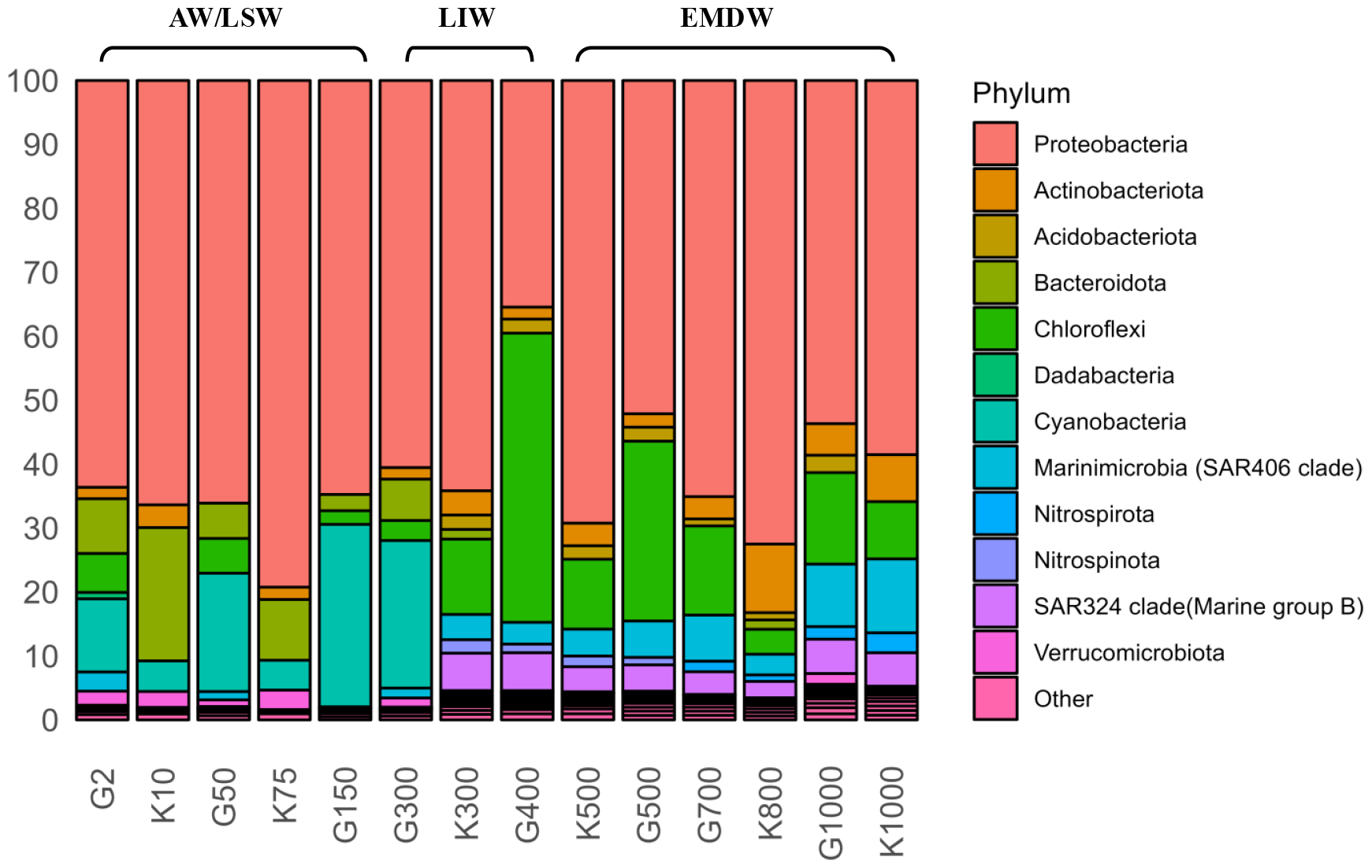
Relative abundance of bacterial taxa at the Phylum level. The three water masses are annotated on top of the barplot (AW/LSW: Surface Water, LIW: Intermediate Water, EMDW: Deep Water). Rare taxa below 1% are classified as “Other”

### Eukaryotic community structure

Unicellular eukaryotic communities differed with both water mass and sampling location (season) (Fig. 5). *Prymnesiophyceae* (Haptophyta) comprised about 35% of total ASVs in Gavdos surface microbial community (G2, winter), five times higher than in Koufonisi (K10, summer). Similar fluctuations occurred in other taxa (*Dinophyceae, Syndiniales*, and *Ascomycota*) between the two AW/LSW samples, with *Syndiniales* being fourfold more abundant in Gavdos (winter) while Fungi (*Ascomycota*) and *Dinophyceae* were higher in Koufonisi (summer). Microbial composition in LIW-G300 sample was more similar to the AW/LSW than the other LIW samples. Radiolaria (*Acantharea* and *Polycystinea*) were favoured in intermediate waters and decreased with depth in contrast with *RAD-B* members of Radiolaria which preferred deeper waters and comprised ∼13% of ASVs at 1000m (Fig. 5). Moreover, the relative abundance of *Syndiniales* and deep-sea acclimated Fungi was elevated in the deepest EMDW samples (G700, K1000).

**Fig. 5.**
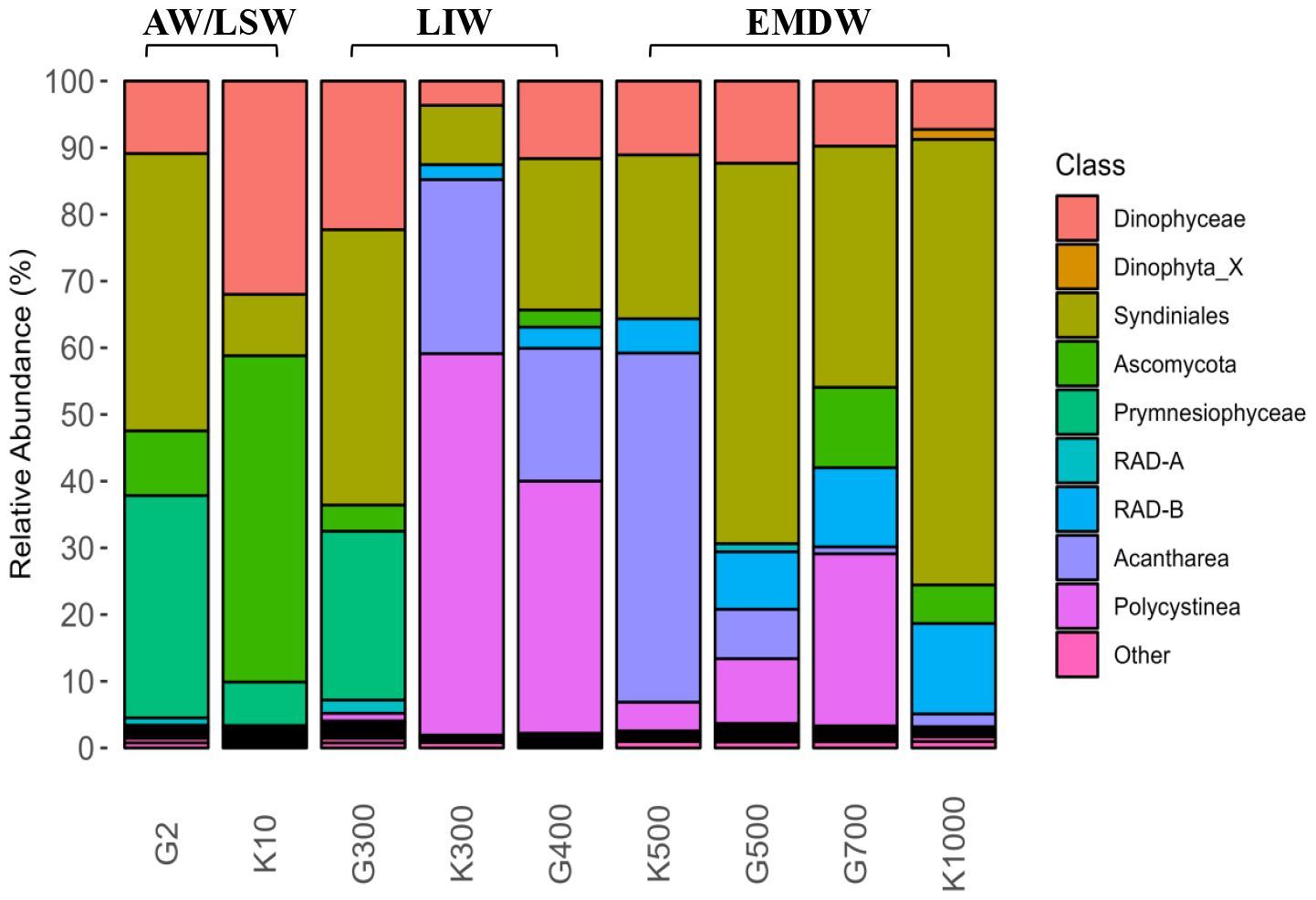
Relative abundance of eukaryotic taxa at the Class level. The three water masses are annotated on top of the barplot (AW/LSW: Surface Water, LIW: Levantine Intermediate Watermass, EMDW: Eastern Mediterranean Watermass). Rare taxa below 1% are classified as “Other”

### Network analysis

The surface AW/LSW co-occurrence network had approximately five times more associations (edges) than the deep EMDW, despite that the latter comprised more ASVs (nodes) (Fig. 6-7, Table S3). Both networks exhibited more positive than negative correlations (Table S4). In the AW/LSW network, Dinoflagellata and Haptophyta were positively correlated with most members of Bacteroidota and Proteobacteria while Fungi (*Ascomycota*, excluding *Sarocladium*) had exclusively positive correlations with Cyanobacteria, Actinobacteriota, and Verrucomicrobiota (Fig. 6). Additionally, Cyanobacteria were negatively related with eukaryotic Haptophytes and Dinoflagellates and were positively associated with Proteobacteria. In EMDW network, approximately 30% of all edges were between Radiolaria and Dinoflagellata, with ∼60% being positive (Fig 7). Deep-sea fungal ASVs were positively correlated with Dinoflagellata in contrast to the AW/LSW network while Radiolaria exhibited exclusively negative connections with Proteobacteria, Nitrospinota (*LS-NOB*), and Picozoa, but positive relationships with Cercozoa and Fungi. Nitrospira was only associated with *RAD-B* taxa of Radiolaria, while Dinoflagellata and Actinobacteriota were negatively correlated, mirroring the surface water mass.

**Fig. 6.**
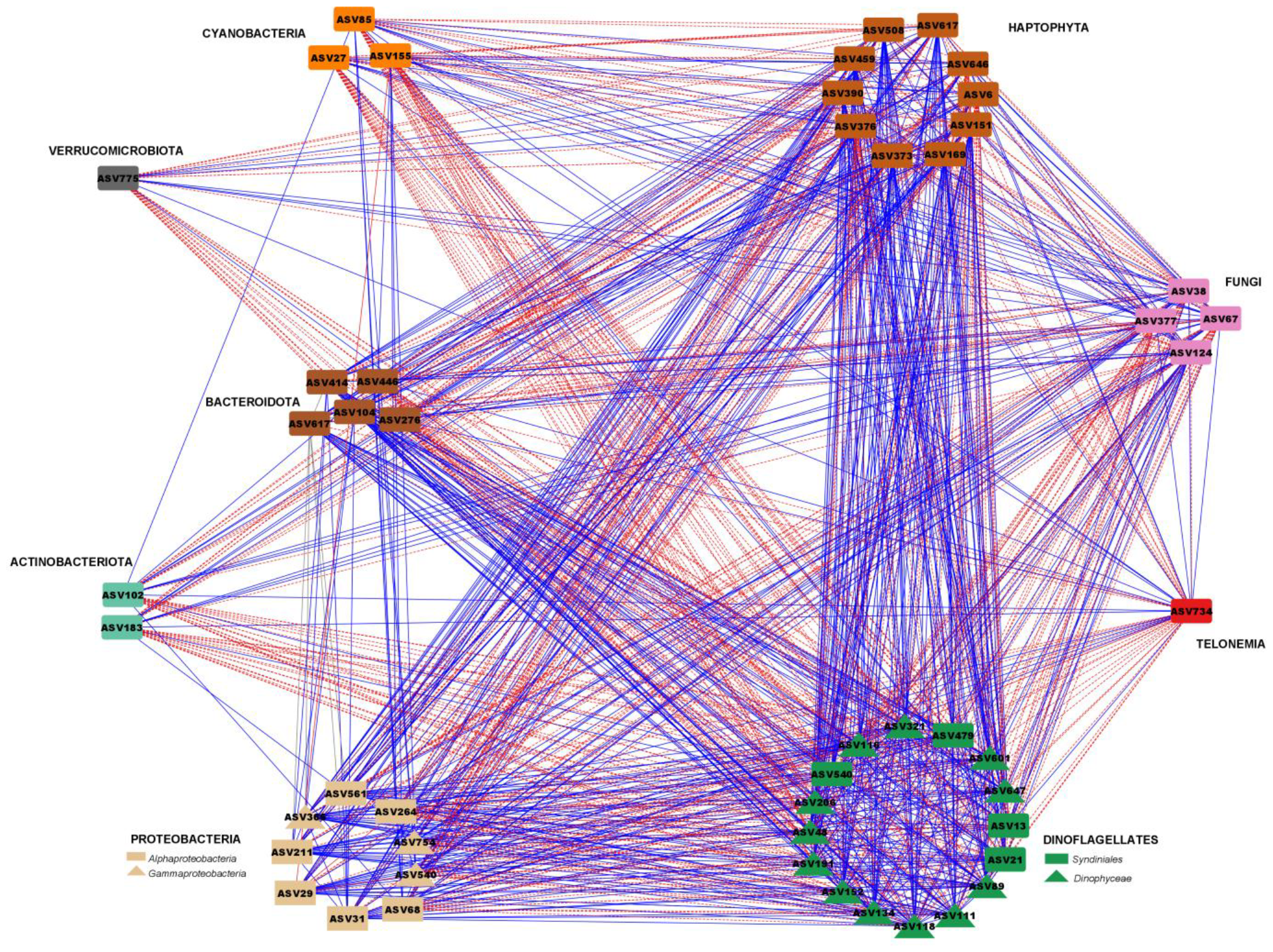
Co-occurrence network diagram of Spearman’s correlations (edges) between the bacterial and unicellular eukaryotic ASVs, (nodes) of the surface water mass. The edges present the statistically significant (*p*<0.05) correlations between ASVs, based on their relative abundance. Blue and red color indicates positive and negative associations respectively

**Fig. 7.**
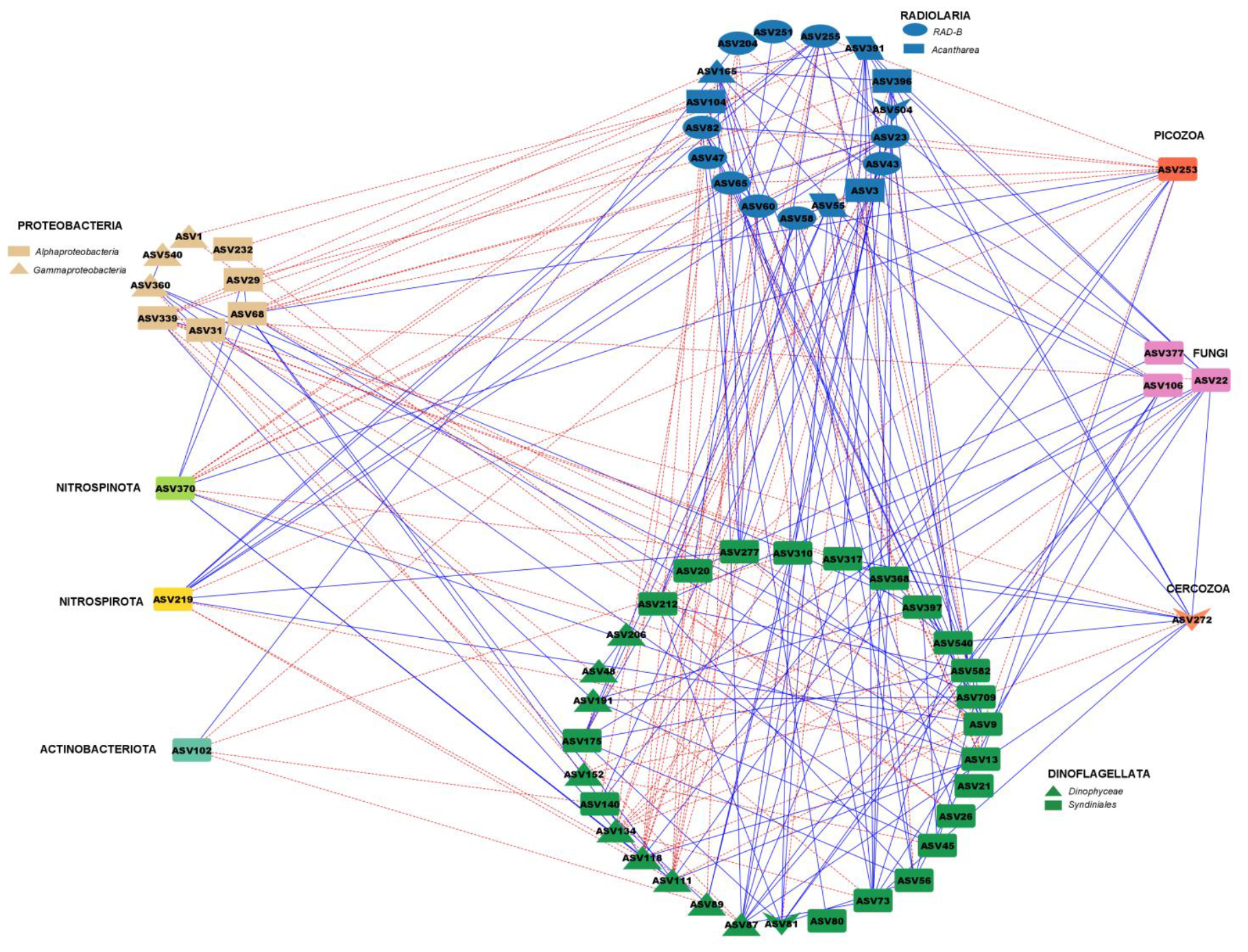
Co-occurrence network diagram of Spearman’s correlations (edges) between the bacterial and eukaryotic ASVs, (nodes) of the deep water mass. The edges present the statistically significant (*p*<0.05) correlations between ASVs, based on their relative abundance. Blue and red color indicates positive and negative associations respectively

## Discussion

This study investigated microbiota structure and relationships between bacterial and unicellular eukaryotic microorganisms in the western part of the Levantine Sea (Cretan Passage) of the poorly investigated Eastern Mediterranean Sea (EMS). Depth-profiled seawater samples were collected from two different stations located within the Hellenic EEZ, approximately 183 km apart, belonging to the same marine ecoregion [33]. As a result, variations in microbial community composition were thought to be attributed on seasonal differences rather than spatial heterogeneity. High-throughput sequencing (HTS) unravelled the bacterial and unicellular eukaryotic diversity and microbial associations were assessed at surface and deep-water layers.

Our findings confirmed that the EMS maintains well-oxygenated conditions throughout its water column with stratified water masses in Koufonisi (summer). However, even though in Gavdos (winter), vertical convection in late February led to water mixing between the surface water (AW/LSW) and intermediate water (LIW) masses down to depths of 300 m, yet uniformal deep (EMDW) waters were observed in both stations (Fig. S1, Fig. S2).

### Microbial community distribution and diversity

The distribution of bacterial and unicellular eukaryotic communities was linked to the seawater layer of origin. Seasonal effects were evident in AW/LSW communities, but the limited number of samples prevented significant testing. Induced vertical mixing in Gavdos (winter) station caused the LIW-G300 sample to exhibit more environmental and biological similarities to the surface than the intermediate water layer. Microbial communities collected from 500m depth clustered with LIW samples, indicating a deeper LIW stratification in the central EMS basin [5, 7]. Sampling location was insignificant for the overall bacterial diversity however, single-cell eukaryotic communities showed significantly higher a-diversity in Gavdos (winter) station in all three water masses (Fig 3D, Fig S5). This was attributed to the lower number of observed species recorded in Koufonisi (summer) in combination with the dominance (lower evenness in the community) of certain eukaryotic groups (Fungi, Dinoflagellates, Radiolaria) probably as a result of feeding and biomass transfer to higher trophic levels in the EMS during this period [34]. Furthermore, the deepest EMDW water mass displayed the highest diversity in both microbial datasets in this part of the EMS, in contrast with studies in the easternmost Levantine region where LIW layer exhibited higher Shannon values [15].

### Photosynthetic community composition and associations in the western Levantine basin (Cretan passage)

Unlike other oceans, highest primary production in the EMS occurs in winter however, the oligotrophic nature of this habitat favours the dominance of picophytoplankton and blooms do not occur [10, 35]. The bacterial fraction of picoplankton, Cyanobacteria, were identified in our AW/LSW samples, with *Synechococcus* consistently dominant, especially in Gavdos (winter) when primary production is higher [10]. *Prochlorococcus* and *Cyanobium* were present only in Gavdos station, in lower abundance than *Synechococcus*, yet they preferred lower AW/LSW depths confirming that, despite their close relation, picocyanobacteria differ ecologically and occupy diverse niches [36, 37]. In contrast to other EMS studies, where seasonal and depth-related variations were observed in the distribution of Cyanobacteria species, in our samples, *Prochlorococcus* was never the dominant Cyanobacteria [15, 18]. In our surface co-occurrence network, Cyanobacteria were exclusively negatively correlated with the mixotrophic *Dinophyceae*, previously associated with phytoplankton grazing [38]. Moreover, *Syndiniales*, the most diverse parasitic group responsible for top-down regulation of *Gyrodinium* (*Dinophyceae*), exhibited a positive relationship with Cyanobacteria confirming that their parasitism is mainly on other marine protists [39]. The bacterial picoplankton was also correlated with haptophytes (*Prymnesiophyceae*) which have been linked with mixotrophy in oligotrophic waters accounting for up to 40% of bacterial grazing [40]. It is worth noting that, unassigned members of *Prymnesiophyceae* (ASV376, ASV646) and *Gephyrocapsa* (ASV6) were the sole representatives of Haptophytes showing a positive association with Cyanobacteria. In contrast, other less prevalent *Prymnesiophyceae* such as *Phaeocystis* (ASV169), *Prymnesium* (ASV373), *Algirosphaera (*ASV459) and *Chrysochromulina* (ASV151) exhibited negative connections, likely attributable to their distinct ecological roles. For instance, *Chrysochromulina*, has been previously linked to increased bacterivory rates, indicating the importance of heterotrophy for its growth, whereas *Gephyrocapsa* may rely more on photoautotrophy than phagotrophy [41].

### Microbial composition and networking in the surface waters of the western Levantine basin (Cretan Passage)

In line with other studies, the heterotrophic SAR11 (*Alphaproteobacteria*) dominated across samples, confirming its high prevalence in oligotrophic environments [42]. Members of SAR11 occupied diverse niches, with *Clade Ia* (ASV29) increasing in summer and *Clade Ib* (ASV68) being more prevalent in winter, influencing their relationships with Haptophytes and Dinoflagellates [17]. Other *Alphaproteobacteria* (*Erythrobacter, OM75*) and *Gammaproteobacteria* (*OM60, SAR86*, and *SAR92*) presented similar seasonal patterns. Despite their common niche for warmer waters, *OM60* (ASV366) and *SAR92* (ASV754) interacted with different eukaryotic groups in our samples. For example, *OM60* was positively associated with phagotrophic Haptophytes, *Dinophyceae* and *Syndiniales*, while *SAR92* was correlated with *Ascomycota* (excluding *Sarocladium*). Moreover, *Pseudohongiella*, another γ-proteobacterium, preferred colder waters and positively interacted with *Prochlorococcus* and *Cyanobium*. Furthermore, the major Bacteroidota, *NS4* and *NS5 marine groups*, were more abundant in Koufonisi (summer) following the picoplankton bloom in winter, confirming their association with the degradation of high molecular weight organic matter (OM) [43]. Marine mycobiome diversity and ecology is generally understudied however they seem to be involved in OM decomposition, nutrient metabolism and parasitism on picoeukaryotic algae [44]. In our samples, marine fungi (*Ascomycota*), mainly represented by *Aspergillus*, were prominent in Koufonisi (summer) AW/LSW sample yet they were not among the significant ASVs in our surface network. However, the negative edges between fungal ASVs, dinoflagellates and haptophytes is in accordance with studies suggesting that marine fungi commonly infect these protists [45].

### Microbial composition and networking in the deeper waters of the EMS basin (Cretan Passage)

The deeper waters of the EMS exhibited high microbial diversity yet associations between species were limited, as evidenced from the fewer number of edges compared to the AW/LSW microbiota network. Below 150 m, taxa linked to nitrogen cycling (*Nitrospinaceae, Chloroflexi, SAR324, SAR406, Nitrospiraceae*) were identified, indicating the significance of nitrogen cycling process taking place here, in alignment with previous EMS studies [15]. *Chloroflexi*, previously associated with recalcitrant OM degradation as well as with deep-sea carbon and sulphur cycling, was particularly abundant in winter LIW waters where OM deposition is higher due to vertical mixing [46].

Prokaryotes like the chemolithoautotroph *Microtrichales* (Actinobacteriota), comammox *Nitrospira*, the sulfur-oxidizing *SUP05 cluster* and members of the microbial dark matter *Marinimicrobia* (*SAR406*) were most abundant in the deepest water layers. ASVs of Dinoflagellates and Radiolaria that often dominate microzooplankton communities, comprised the majority of nodes in our EMDW network [4]. In particular, the Radiolaria species of *Acantharea* and *Polycystinea* dominated the LIW and upper EMDW while unclassified *RAD-B* taxa were increased in deeper waters. In oligotrophic habitats, like the EMS, Radiolaria are considered as important players in the biological carbon pump even thought their trophic role is unclear [47]. However, they have been categorized as non-constitutive mixotrophs ingesting specific algal prey (eSNCM) like Dinoflagellates and Haptophytes [48]. This ecological role is in agreement with our findings explaining the high number of positive edges between Dinoflagellata and Radiolaria. In our findings, Nitrospinota and Proteobacteria seem to act as prey for Radiolaria based on the exclusively negative associations between them. Moreover, the uncultivated *Syndiniales*, which comprised 60% of the total abundance in the deepest samples, are considered a black box in protistology [49]. Dinoflagellates seem to act as predators for the prokaryotic *Nitrospira* and *Microtrichales* based on our results while deep-sea fungal taxa were also observed (*Aspergillus, Lecanicillum,Cladosporium*) in accordance with previous studies [50]. Finally, positive relationships of deep-sea fungi with Radiolaria and Dinoflagellates might suggest a symbiotic-mutualistic or commensalistic relationship between these species.

Overall, our study aims to investigate that in the oligotrophic EMS basin, microbial communities can be differentially shaped according to distinct water masses. Through analysing samples from two sampling stations in this highly oligotrophic oceanic region, we observed distinct bacterial and unicellular eukaryotic communities with water mass. The eukaryotic community appeared to be more diverse throughout the water column, particularly in LIW layer, at Gavdos (winter) station than in Koufonisi (summer). However, more interseasonal sampling expeditions are required for this to be statistically verified. The microbial network analysis revealed interspecies associations, indicating the increased number of ecological niches present in the surface layer compared to the deeper water mass. These findings underline the necessity for further exploration and understanding of microbial ecology and associations in one of the world’s most understudied marine habitats, especially if we consider the heavy anthropogenic stress and the prediction of climate change impact in this basin.

## Notes

### Competing Interest Statement

The authors have declared no competing interest.

