## Supplementary Material for "Distinct communities of Bacteria and unicellular eukaryotes in the different water masses of Cretan Passage water column (Eastern Mediterranean Sea)"

**Supplementary Materials:**

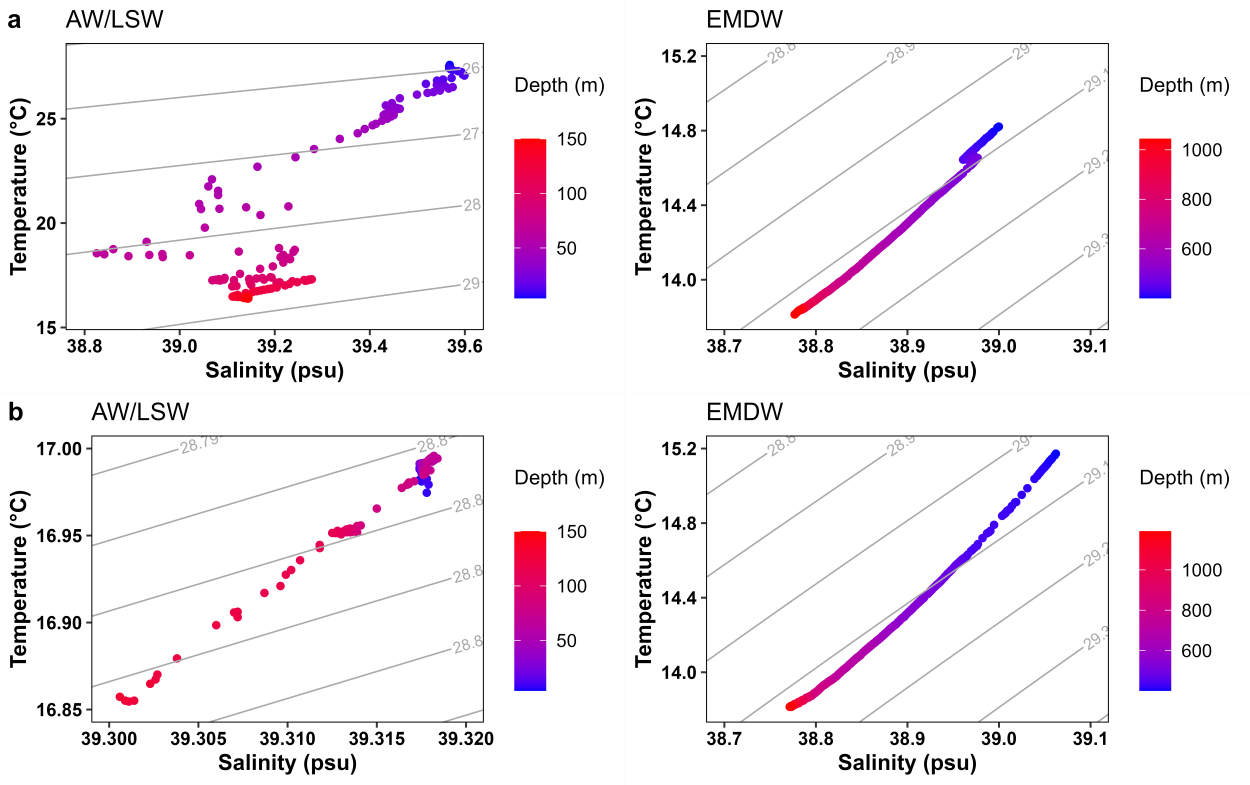

**Fig. S1** Temperature-Salinity diagrams for surface (AW/LSW) and deep (EMDW) water masses of a) Koufonisi and b) Gavdos station. Thin grey lines represent isopycnals of density in kg m^-3^

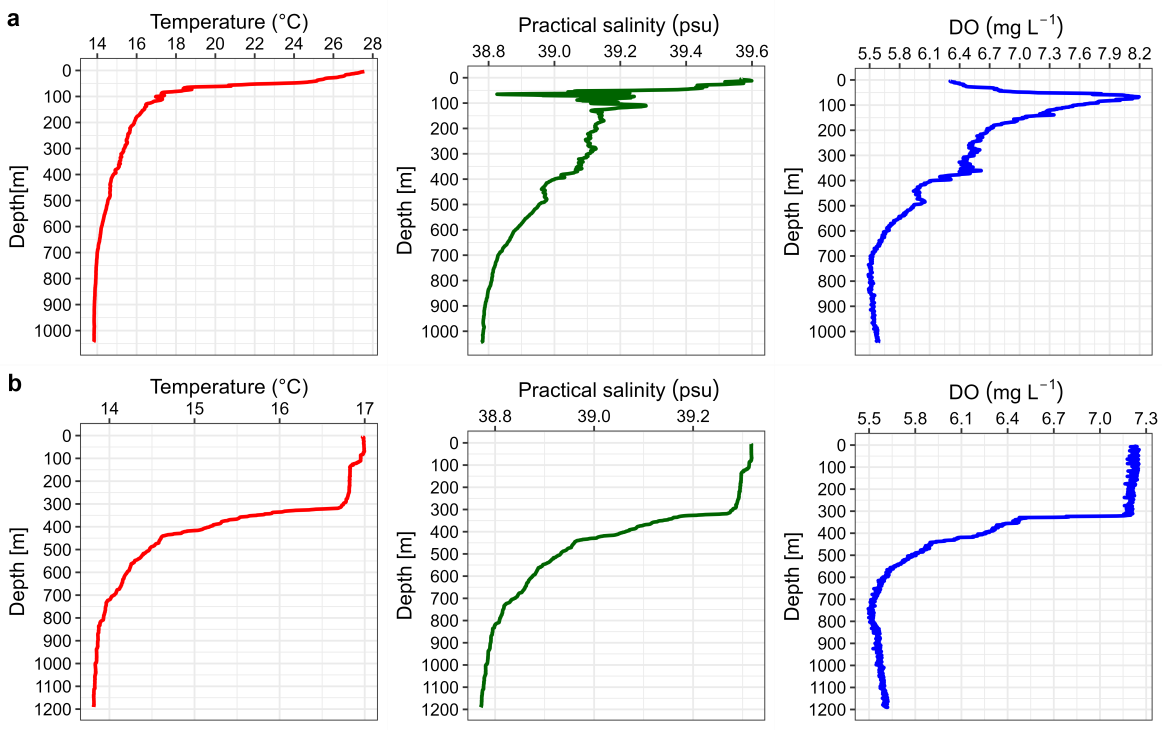

**Fig. S2** Temperature, salinity and dissolved oxygen profiles for a) Koufonisi and b) Gavdos station

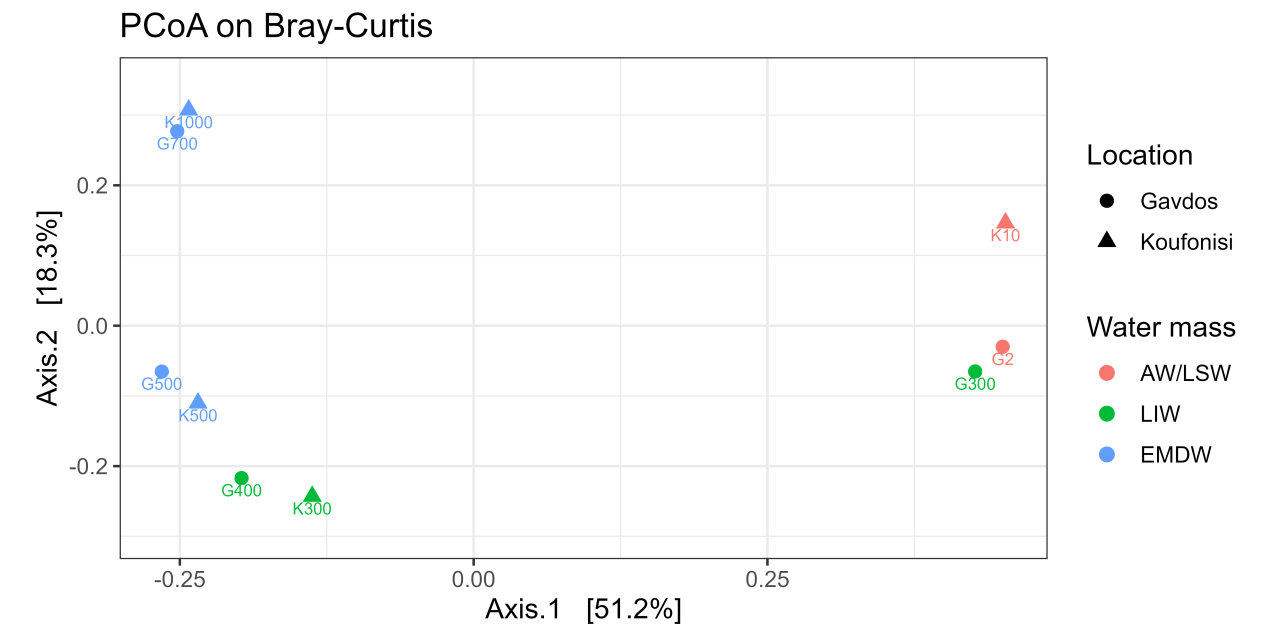

**Fig. S3** Principal coordinate analysis of unicellular eukaryotic samples based on Bray-Curtis dissimilarity distances

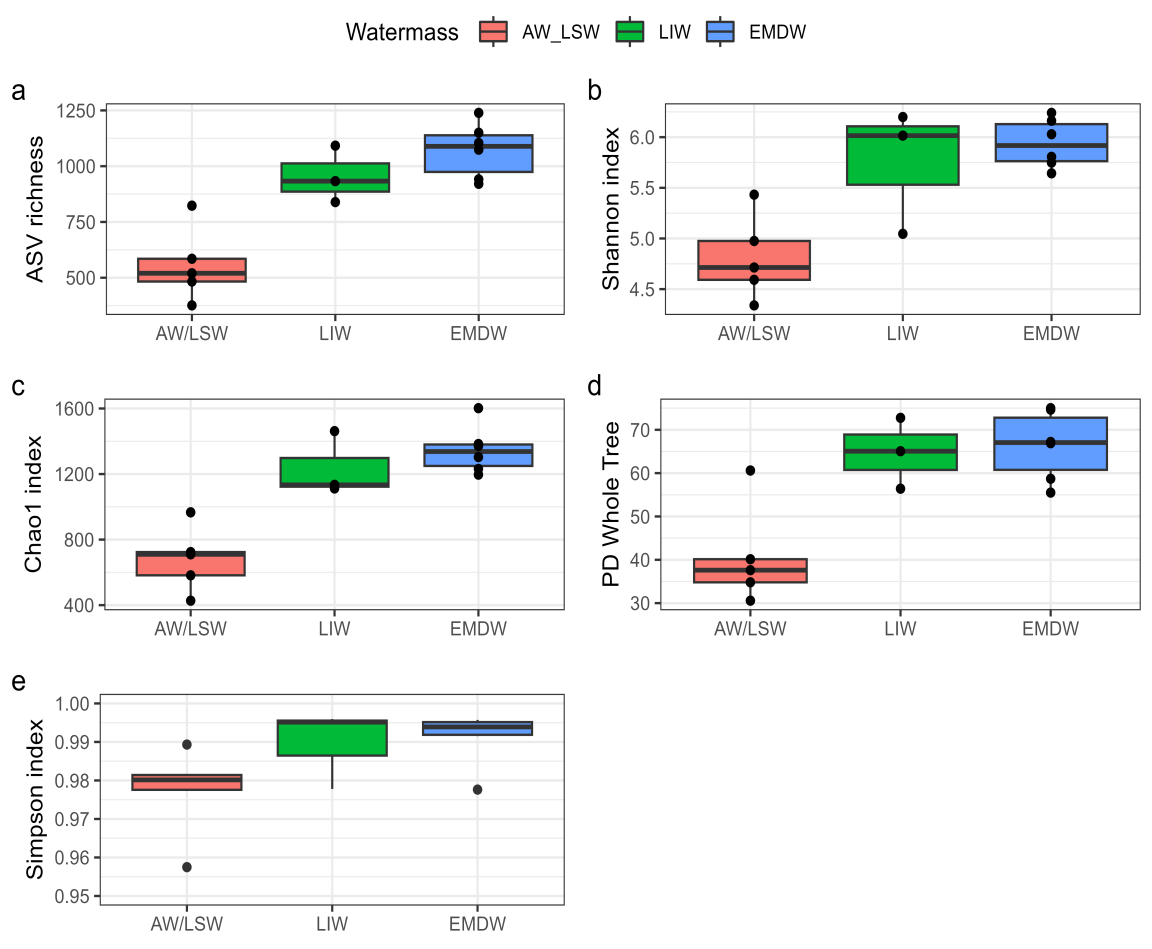

**Fig. S4** Boxplot of alpha diversity indices of bacterial samples for each water mass

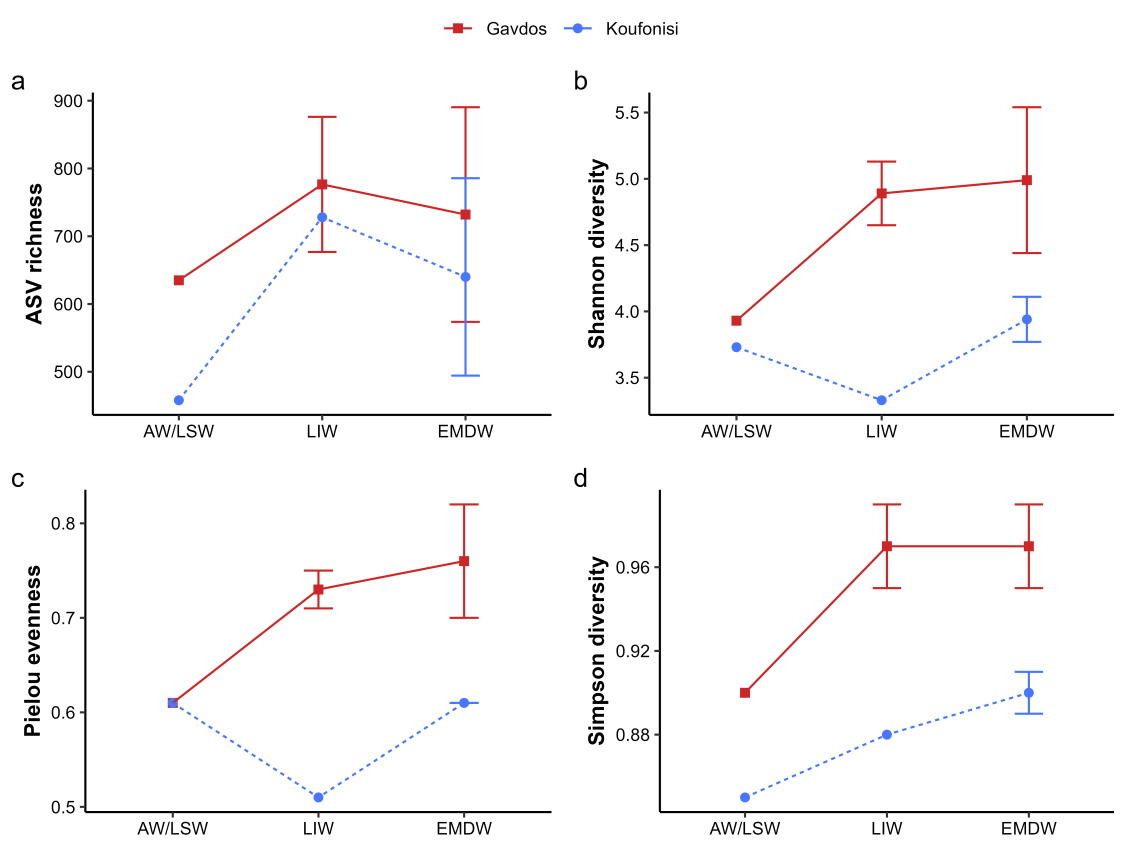

**Fig. S5** Alpha-diversity metrics of unicellular eukaryotes in each water mass for every sampling station

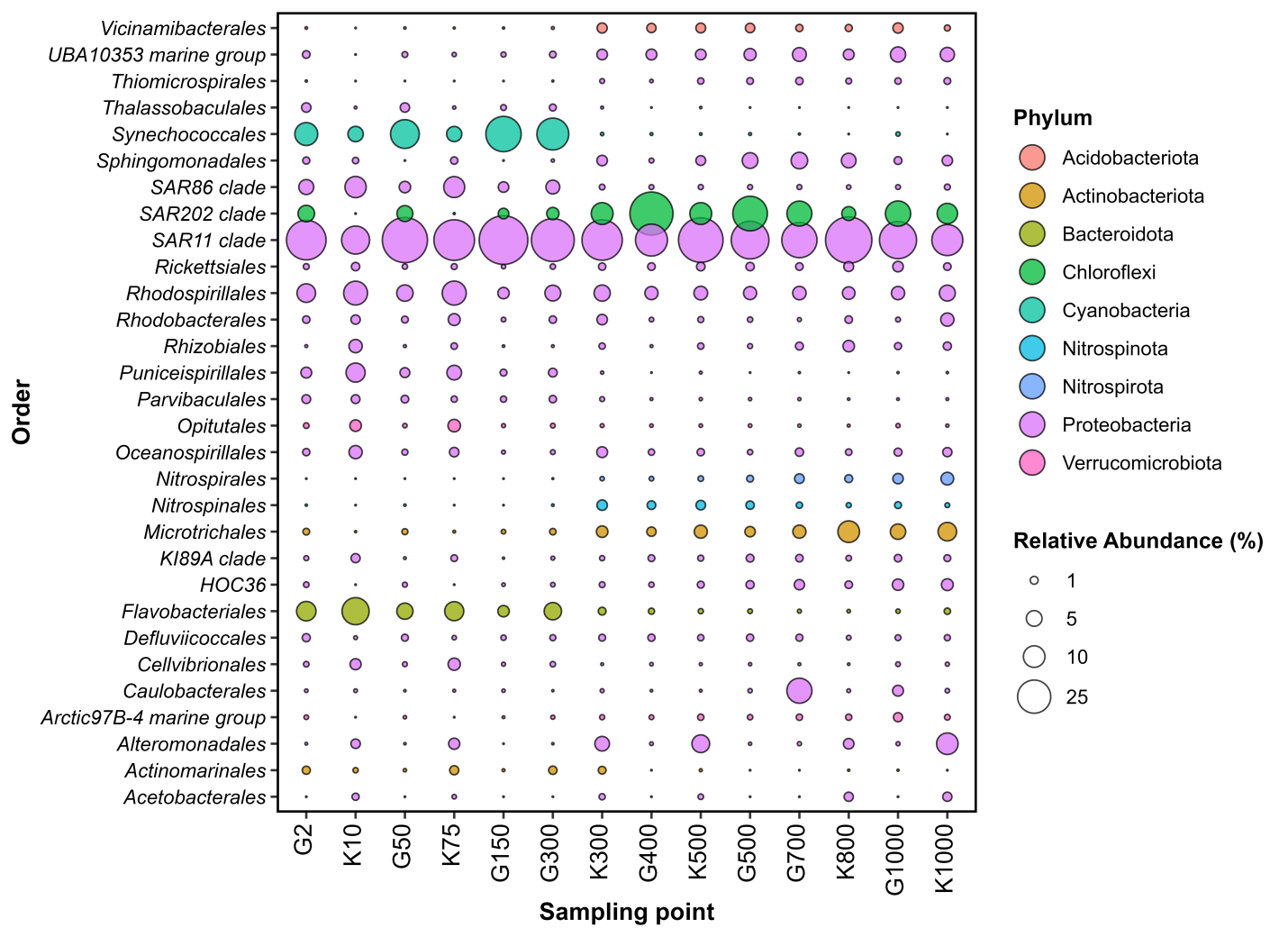

**Fig. S6** Bubbleplot of relative abundances of bacterial taxa at the *Order* level

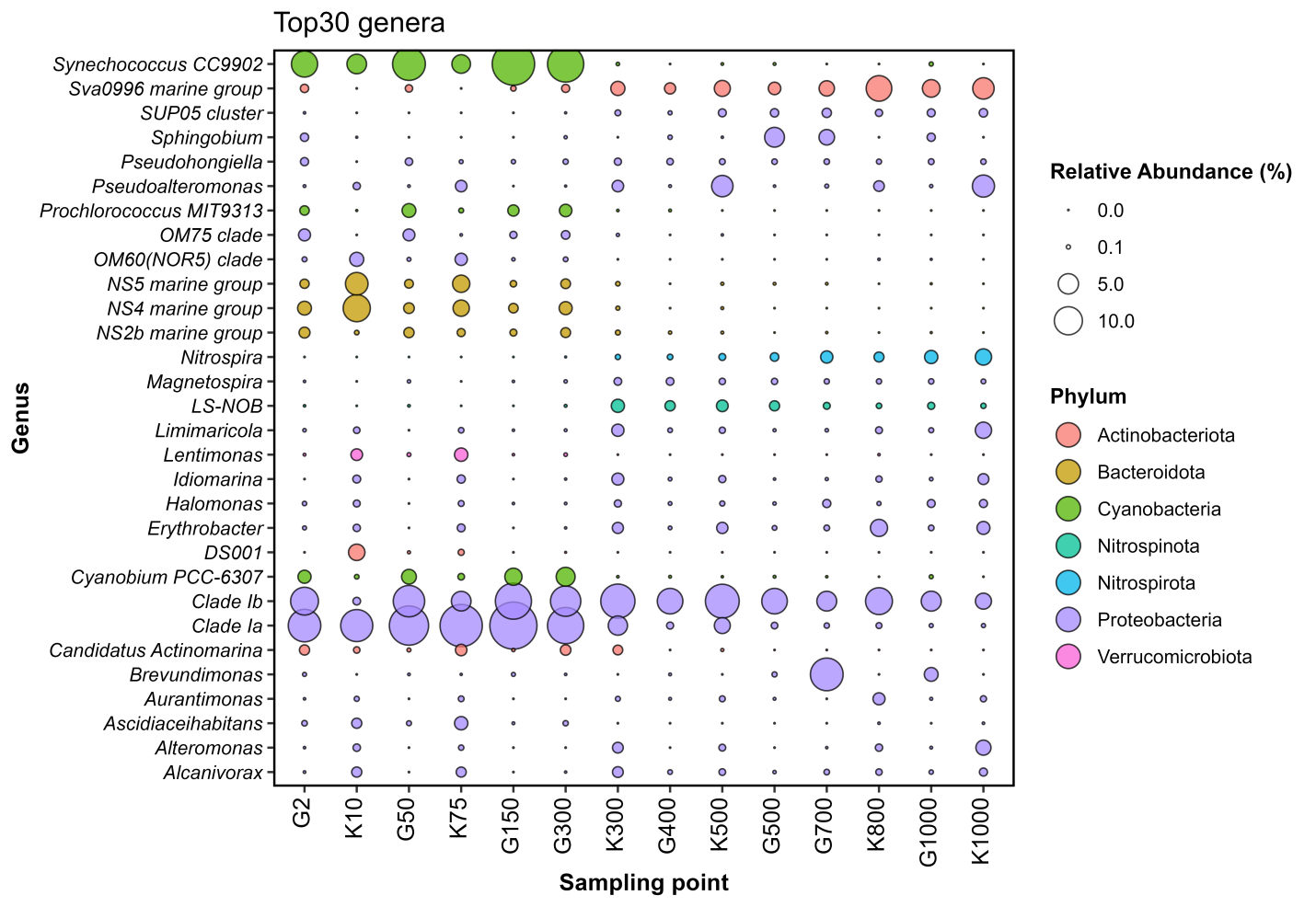

**Fig. S7** Bubbleplot of relative abundances of Top30 bacterial genera

**Table S1** ANOVA and Tukey HSD significance test of differences in alpha-diversity of bacterial communities between the three water masses.

|  |  | ANOVA | |  |  | Tukey HSD test | | |
| --- | --- | --- | --- | --- | --- | --- | --- | --- |
| Diversity metric |  | ***p*-value** | **F-value** |  | | **AW/LSW-LIW** | **LIW-EMDW** | **AW/LSW-EMDW** |
| Shannon |  | 0.00189 | 11.69 |  | | 0.01994 | 0.79 | 0.00181 |
| Simpson |  | 0.0777 | 3.25 |  | | 0.22189 | 0.97 | 0.07825 |
| PD whole tree |  | 0.00226 | 11.14 |  | | 0.01382 | 0.97 | 0.00264 |

**Table S2** ANOVA significance results of differences in alpha-diversity of unicellular eukaryotic communities between the two sampling locations (season) and the three water masses.

|  |  | Location (season) | |  | Watermass | |
| --- | --- | --- | --- | --- | --- | --- |
| Diversity metric |  | ***p*-value** | **F-value** |  | ***p*-value** | **F-value** |
| Shannon |  | 0.018 | 12.11 |  | 0.300 | 1.54 |
| Simpson |  | 0.001 | 46.92 |  | 0.012 | 12.14 |
| PD whole tree |  | 0.212 | 2.041 |  | 0.121 | 3.31 |

**Table S3** Characteristic parameters of surface and deep co-occurrence networks.

| **Network** | **SURFACE** | **DEEP** |
| --- | --- | --- |
| **Number of nodes** | 51 | 63 |
| **Number of edges** | 1122 | 237 |
| **Positive correlation (%)** | 53 | 61 |
| **Avg. number of neighbors** | 44 | 11.5 |
| **Network diameter** | 2 | 4 |
| **Network radius** | 1 | 2 |
| **Characteristic path length** | 1.12 | 1.35 |
| **Clustering coefficient** | 0.923 | 0.861 |
| **Network density** | 0.88 | 0.767 |
| **Network heterogenity** | 0.173 | 0.317 |
| **Network centralization** | 0.125 | 0.114 |
| **Connected components** | 1 | 9 |
